# Timed sequence task: a new paradigm to study motor learning and flexibility in mice

**DOI:** 10.1101/2023.06.30.547172

**Authors:** Anna Urushadze, Milan Janicek, Alice Abbondanza, Helena Janickova

## Abstract

Motor learning and flexibility allow animals to perform routine actions efficiently while keeping them flexible. There is a number of paradigms used to test cognitive flexibility but not many of them focus specifically on learning of complex motor sequences and their flexibility. While many tests use operant or touchscreen boxes that offer high throughput and reproducibility, the motor actions themselves are mostly simple presses of a designated lever. To focus more on motor actions during the operant task and to probe the flexibility of these well-trained actions, we developed a new operant paradigm for mice, the, “timed sequence task”. The task requires mice to learn a sequence of lever presses that have to be emitted in precisely defined time limits. After training, the required pressing sequence and/or timing of individual presses is modified to test the ability of mice to alter their previously trained motor actions. We provide a code for the new protocol that can be used and adapted to common types of operant boxes. In addition, we provide a set of scripts that allow automatic extraction and analysis of numerous parameters recorded during each session. We demonstrate that the analysis of multiple performance parameters is necessary for detailed insight into animals’ behavior during the task. We validate our paradigm in an experiment using the valproate model of autism as a model of cognitive inflexibility. We show that the valproate mice show superior performance at specific stages of the task, paradoxically due to their propensity to more stereotypic behavior.

**Significance Statement:** Cognitive flexibility impairment is a crucial component of many neurological disorders and it is frequently evaluated in animal models. As the commonly used tests usually do not focus on motor learning and the ability to adapt motor sequences, we designed a new paradigm to evaluate motor learning and its flexibility. The timed sequence task is automatized and easily accessible as it is based on widely available operant boxes. During the training, the task requires precise timing of each action to force stereotypic performance. Its relative complexity allows detailed analysis of multiple parameters and therefore detailed insight into animal’s behavior. The task can be used to reveal and understand subtle differences in motor and operant learning and flexibility.

**Graphical Abstract:** 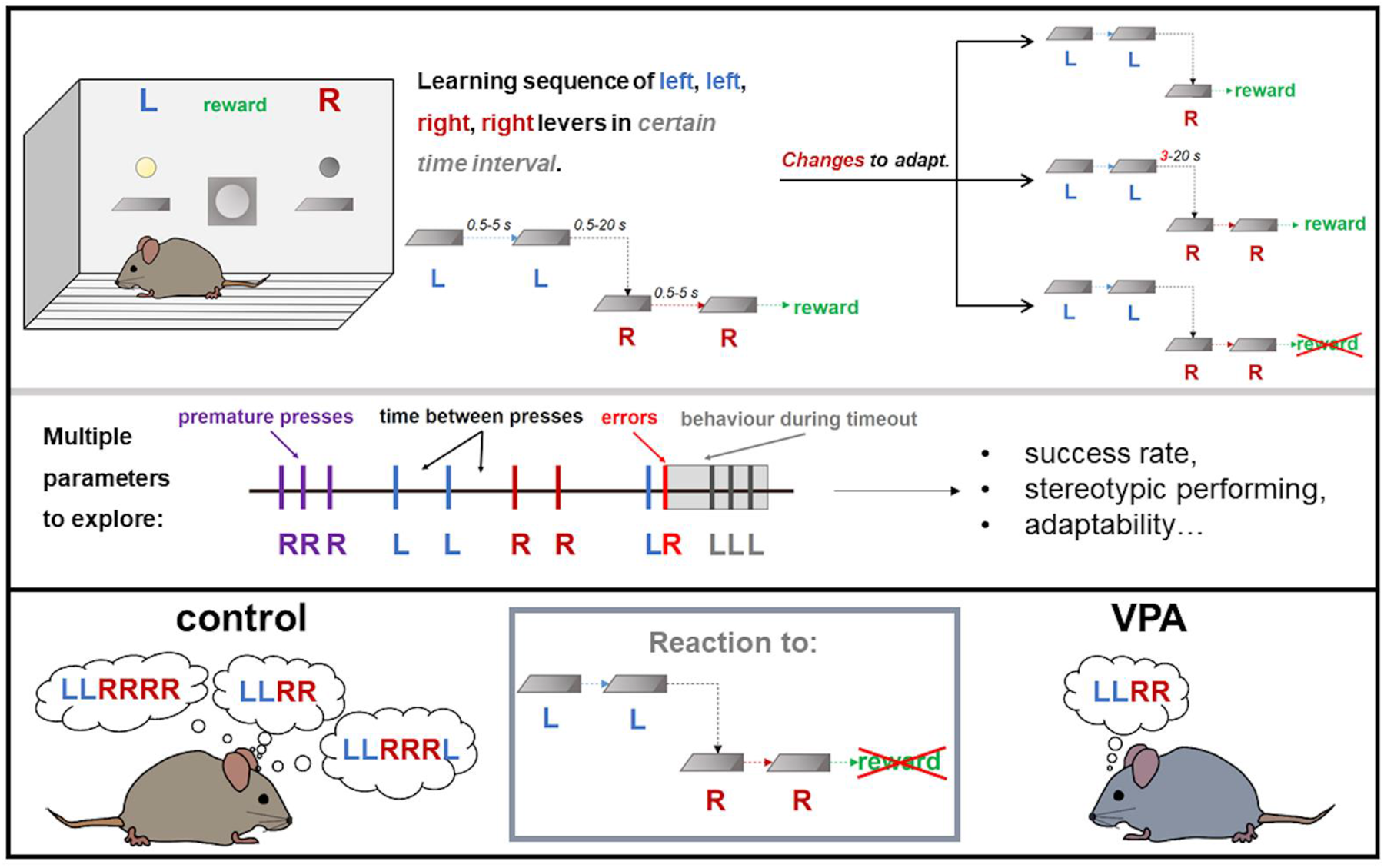

## Introduction

Acquisition of new motor actions and their successful automatization is a crucial part of animals’ behavior. The animals have to be able to not only learn the motor action but also flexibly change it when required. Various tasks for testing of cognitive flexibility are performed in conventional operant or in touchscreen-based boxes (Mar et al., 2013). Alternatively, a variety of maze-based tasks (Vorhees and Williams, 2006) can be used. In most of these paradigms, the original motor action is relatively simple (a lever press or a nose poke) and in the flexibility phase, it is transformed into its mirror image. In these paradigms, it may be difficult to capture subtle changes in animal’s behavior as a simple motor action does not allow for a detailed insight (Keeler et al., 2014). In literature, several paradigms can be found that are suitable for testing more complex motor learning and automatization (Keeler et al., 2014; Geddes et al., 2018; Dhawale et al., 2021; Janickova et al., 2021; Turner et al., 2022; Wolff et al., 2022). However, these tasks usually do not involve flexibility part and, more importantly, they often require a specialized equipment and/or technical expertise which makes them less accessible. Notably, some of these more complex tasks have been so far performed only with rats and their adaptability to mice may not prove easy.

To overcome the aforementioned difficulties, we created a timed sequence task where mice perform a heterogeneous sequence of four presses in a highly stereotypic way. To ensure this, mice are taught to perform the individual presses within pre-defined time limits, to make the actions in individual trials highly uniform. Once the complex sequence is well trained, it can be repeatedly modified at different steps, to implement the flexibility component of the task. In addition, the task structure allows to collect and analyze numerous behavioral parameters. The present protocol can be realized with commonly available operant boxes equipped only with two non-retractable levers and a feeder. The protocol code is programmed in a proprietary but common programming language that is used with a common type of operant boxes. The code is open and can be easily modified to create new variations of the task. To shorten the tedious analysis of multiple task parameters, we also provide a set of scripts in Python for automatized processing of the data and for user-friendly manual checking of lengthy task logs if needed. As a first choice, we tested our new protocol in the valproate mouse model of autism (Nicolini and Fahnestock, 2018), a model with a prominent social impairment, impairment of motor skills and a higher propensity to habitual and repetitive behavior (Nicolini and Fahnestock, 2018; Tartaglione et al., 2019). In this model, we show specific alterations in several parameters of the task, together providing evidence of increased habitual performance and slight motor impairment in these animals.

## Materials and Methods

### Animals, preparation and testing of the VPA model

Male mice of C57BL/6J genetic background were used and purchased from Charles River Laboratories. They were group-housed in a room with controlled temperature and 12-hour light cycle. All experimental procedures complied with the directive of the European Community Council on the use of laboratory animals (2010/63/EU) and were approved by the animal care committee of the Czech Academy of Sciences. The behavioral testing started at 2 months of age. Prenatal exposure to valproic acid (VPA) was used to induce autistic phenotype, according to the published protocol (Lucchina and Depino, 2014). Pregnant females were injected subcutaneously at the gestation day 12.5 with VPA sodium salt (600 mg/kg, Sigma) in a volume of 5 ml/kg or, for controls, with vehicle only (saline). In total, we produced 11 control males and 11 males exposed to VPA.

### Confirmation of the autistic phenotype

In 2-month old mice, we confirmed the autistic phenotype (Ergaz et al., 2016) using three tests in the following order: social preference test (Nadler et al., 2004), hole-board test to examine compulsivity (Martos et al., 2017) and sticky tape test to evaluate dexterity (Bouet et al., 2009). The social preference test was performed in a three-chamber apparatus (90 × 23 × 23 cm). A mouse was placed in the central chamber and habituated for 5 minutes. A wire mesh pen cup was allocated in each lateral chamber, with one side containing a juvenile male. During 10-minute test session, the time spent by interacting with the juvenile or with the empty cup was manually scored. For the hole-board test (Martos et al., 2017), mouse was placed in a box (40 × 40 cm) equipped with a double floor containing 16 equidistant holes. We recorded sequences of nose pokes into individual holes during 30 minutes and calculated the probability of returning to the same hole. For the sticky tape test (Bouet et al., 2009), triangle-shaped pieces of tape were attached to the front limbs and the time until the removal of both tapes was measured, with the maximum time 5 minutes. The procedure was repeated 3x for each mouse and averages from the two shortest times were analysed.

### Timed sequence task: overview

The task can be performed in standard operant boxes equipped with fixed levers. In the timed sequence task, mice have to learn a precisely timed sequence of lever presses without any error or interruption. The sequence is acquired in several stages. After successful acquisition, the sequence or the time intervals can be changed in multiple probe sessions to examine the ability to change the previously learned motor program. Each set of probe sessions is followed by baseline sessions to re-establish the original performance. To perform the task, we used a standard operant box (MED-307A-B1, MedAssociates) placed in a sound-attenuating cubicle and operated by Med-PC V software (MedAssociates). The box featured two fixed levers and a feeder located between them, for the automatic delivery of reward pellets. Above each lever, a cue light was located indicating when the lever was active and should be pressed. The correct execution of each trial was rewarded with a chocolate cereal pellet (20 mg, Bio-Serv). The main house light in the box was kept off throughout the training. Mice underwent daily training sessions between 8am-5pm, 5 times per week. The completion of the whole training and testing required approximately 40 training/testing days (8 weeks) for a control mouse. The majority of animals were able to learn the task and complete the whole procedure. However, we defined criteria in Table 1 for each stage to remove non-performing animals from the study. The individual stages of the training and testing are listed in Figure 1 and Table 1 and in details described below. The respective codes for each stage are available at https://doi.org/10.5281/zenodo.7881104.

**Figure 1:**
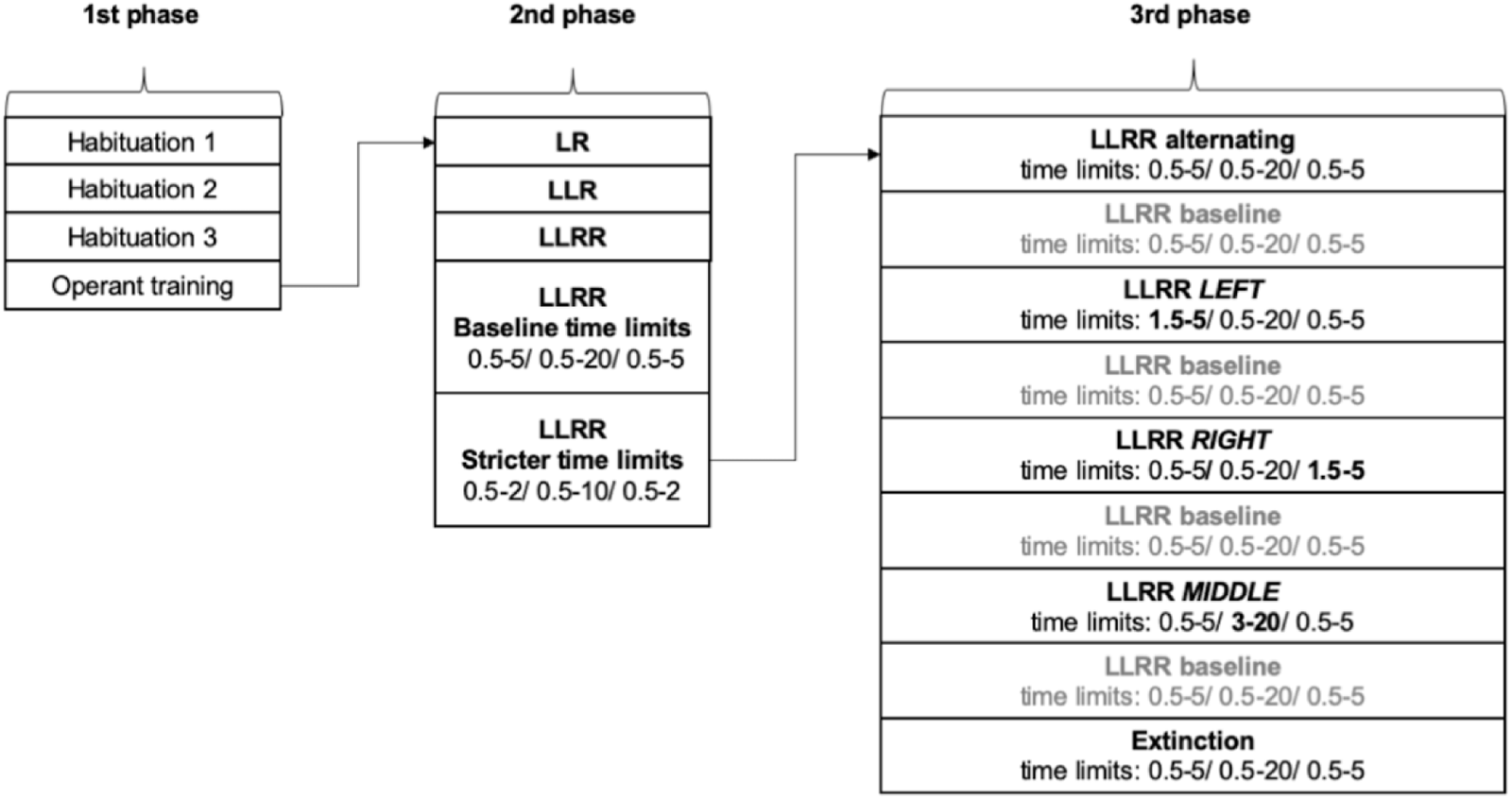
Schematic of the training stages during the timed sequence task. The training consists of 3 main phases: 1/ basic operant training, 2/ acquisition of the timed LLRR sequence and 3/ alterations of the previously learned timed sequence. In the last stage, each alteration is followed by 1-3 sessions of the baseline timed sequence protocol, to ensure mice return to their baseline performance.

**Table 1:** Exclusion criteria used for the individual training stages of the task.

| Training stage | Exclusion criteria |
| --- | --- |
| Habituation 1 | No exclusion criteria |
| Habituation 2 | Mouse does not reach performance criteria within 10 sessions |
| Habituation 3 |  |
| Operant training |  |
| LR sequence | Mouse does not reach performance criteria within 10 sessions |
| LLR sequence |  |
| LLRR sequence |  |
| LLRR with baseline time limits | Mouse does not reach performance criteria within 10 sessions and performs less than 10 trials on the last (10th) session. |
| LLRR with stricter time limits |  |
| LLRR alternating: 3x LLRR + 1x LLR | No exclusion criteria |
| LLRR baseline | Mouse performs less than 10 trials on the last (3rd) baseline session. |
| LLRR LEFT | No exclusion criteria |
| LLRR baseline | Mouse performs less than 10 trials on the last (3rd) baseline session. |
| LLRR RIGHT | No exclusion criteria |
| LLRR baseline | Mouse performs less than 10 trials on the last (3rd) baseline session. |
| LLRR MIDDLE | No exclusion criteria |
| LLRR baseline | Mouse performs less than 10 trials on the last (3rd) baseline session. |
| Extinction | No exclusion criteria |

### Food restriction

For the food motivated testing, two weeks before the training initiation, mice were weighed and the range of 85 – 90 % of their original weight was determined as the target weight. Mice were weighed and fed daily to maintain their target weight throughout the training and testing.

### Timed sequence task: description of individual stages

#### 1st Phase, Operant training

##### Habituation 1

On the first day of training, mice were placed in the operant box with the feeder off for 10 minutes. No reward was placed in the feeder and all mice automatically advanced to the next training step on the next day.

##### Habituation 2

At the beginning of this session, four reward pellets were manually placed into the feeder and mice were left in the box for 10 minutes. At the end of the session the pellets were checked and only the mice that consumed all pellets advanced to the next training stage. Otherwise, the session was repeated.

##### Habituation 3

During last habituation, mice stayed in the box for 30 minutes while the reward was automatically delivered every 2 minutes. Mice had to consume all 15 reward pellets in order to advance to the next stage. Otherwise, the session was repeated.

##### Operant training

After habituation, mice had to associate lever presses with the delivery of the reinforcement. During the initial operant training, every (left and right) lever press was rewarded and the levers became active immediately after the reward delivery. The session ended after mouse earned 40 rewards or after 30 minutes, whichever occurred first. After mice successfully earned 40 rewards within a session, they were moved to the next stage. Otherwise, the session was repeated.

#### 2nd Phase, LLRR Acquisition

##### LR sequence

To avoid any side bias, mice were initially trained to perform a left-right sequence. To obtain a reward, mice had to complete the whole LR sequence with no errors (that is, sequences “LLR” or “RLR” were not rewarded). After every error, a 5s time out (TO) interval was introduced during which the presses were not counted as correct and did not yield any reward. The session ended after 60 minutes or when the mouse earned 40 rewards, whichever occurred first. In order to proceed to the next training stage, mice had to earn 40 rewards within the session per day for three consecutive days. This criterion and TO settings remained the same for all stages of the 2nd Phase. The 40 rewards criterion was selected based on our pilot studies as it was feasible and yet sufficiently challenging for mice.

##### LLR sequence

This stage was similar to the previous with the addition of another left press, forming the sequence LLR (left, left, right).

##### LLRR sequence

The final sequence LLRR (left, left, right, right) consisted of four presses that had to be performed without any errors to be rewarded. Up to this point, intervals between the correct presses were not specified.

##### LLRR with baseline time limits

At this stage, every press had to be emitted within a certain interval after the preceding press. We call the time limits introduced at this stage as “baseline time limits” and this particular timed sequence was used for the re-establishing of the baseline performance throughout the following testing. The example illustration of a correct and incorrect trial at this stage is shown in Figure 2.

The time intervals we used were 0.5-5 s between the 1st and 2nd press (L-L), 0.5-20 s between the 2nd and 3rd press (L-R) and again 0.5-5 s between the 3rd and 4th press (R-R). To obtain the reward, the sequence had to be performed with no errors and interruption, within the pre-defined time limits. The cue lights indicated the activation of the respective lever after the previous press. As in the preceding training stages, the punishment 5 s TO was introduced after every either pressing or timing error.

**Figure 2:**
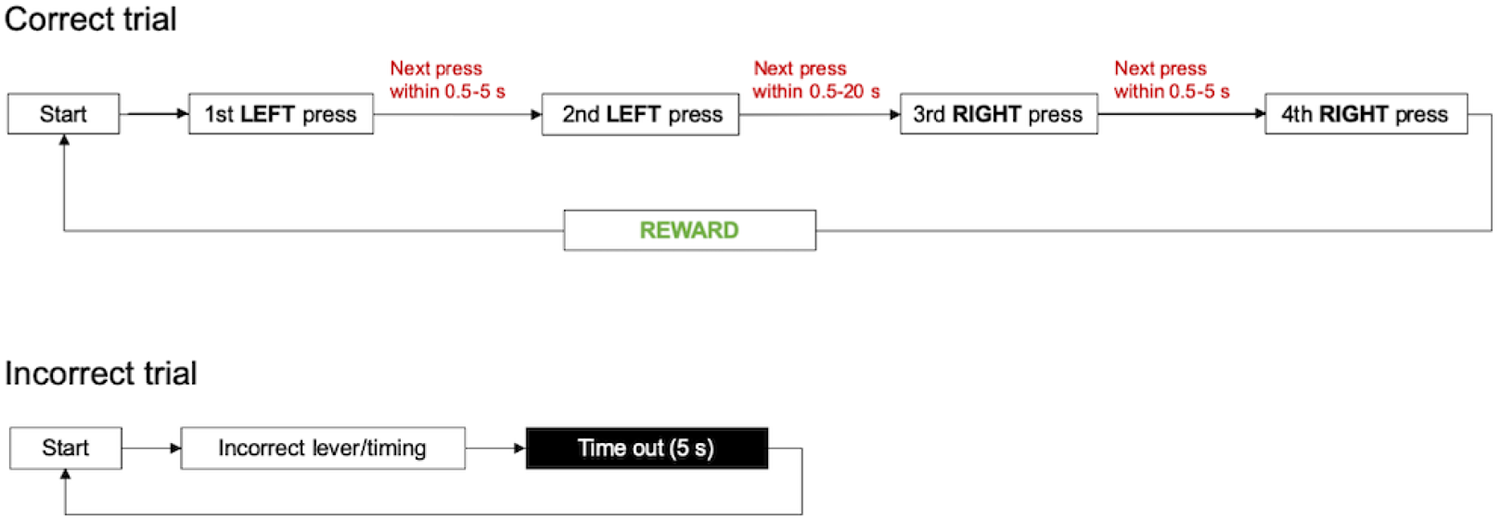
Illustration of the correct and incorrect trial in the baseline timed LLRR sequence. Only complete correct sequences performed within the pre-defined time intervals are rewarded. After each incorrect trial (resulting from an incorrect press or a timing error), 5s timeout is introduced.

##### LLRR with stricter time limits

At this stage, the mice had to perform the sequence within stricter time limits while all other parameters remained the same. The limits were set as 0.5-2 s (L-L), 0.5-10 s (L-R) and 0.5-2 s (R-R). At this stage, some mice started having difficulties to fulfil the criterion. If this was the case, we trained the animals for the maximum of ten days and after that, we moved them automatically to the next stage (also see Table 1 for exclusion criteria).

#### 3rd Phase, LLRR alterations

##### LLRR alternating: 3x LLRR + 1x LLR

After training mice in the baseline timed sequence, we started alternating the pressing sequence and its timing in order to examine the cognitive flexibility of the animals. First, we tested the effect of an alternated sequence by changing the fourth sequence during the session to LLR. Therefore, in every fourth trial the last press had to be omitted and potential execution of the second right press was counted as an error. Hence, the mouse could choose either the earlier reward or the stereotypic completion of the sequence before getting the pellet. The time intervals between the presses were the same as in the baseline LLRR sequence. No performance criteria were set from this stage onwards and all mice were tested in the alternated sequence for three consecutive days. After that, they were run in 1-3 baseline session (LLRR sequence with baseline limits) until they earned 40 rewards at baseline or for the maximum of 3 days.

##### LLRR LEFT

After re-establishing the baseline performance, we started alternating the time limits for individual presses. First, we changed the interval between the 1st and 2nd press (L-L) to 1.5-5 s, forcing the mice to wait for additional 1 s before the second press. The other intervals between presses remained unchanged (0.5-20 s and 0.5-5 s, respectively).

##### LLRR RIGHT

This time, we changed the last interval between the 3rd and 4th press (R-R) to 1.5-5 s.

##### LLRR MIDDLE

Finally, we changed the interval between the two middle presses (L-R) to 3-20 s. After three days of testing, mice were moved to their last baseline session(s) and then to the final stage.

##### Extinction

On the last day of our paradigm, we submitted mice to the extinction session with no reinforcement. The correct sequence was identical to the LLRR sequence with baseline time limits, including the cue lights and the 5 s TO after incorrect trials, however, no reward was delivered after completing a correct sequence.

### Timed sequence task: data analysis

The boxes were operated by the MedPC-V software that was also used to record all events that were defined in the task code. The resulting task log shows all successive events in a row (one row per session) as decimal numbers: the number before decimal point indicates the time of the respective event that elapsed from the last recorded event. The number after the decimal point indicates the specific event using a respective code as defined in the task. The task events recorded in the log included individual lever presses, both correct and incorrect and presses during the TO interval, and reward delivery after the completion of correct sequences. Main summary parameters for each session (e.g. total number of presses) were recorded in a text file generated for each session. For more detailed analysis, we generated a comprehensive task log containing all sessions of the study and analyzed various performance parameters using the log and the analysis scripts created for that purpose. In addition to the individual analysis scripts, we also created a script allowing an easy and quick searching in the extensive task logs and visualization of defined time points during the session. The complete task log containing data from this study can be found at https://doi.org/10.5281/zenodo.7875357. Summary of all task codes and parameters we used is shown in Tables 2-4. The codes and the analysis scripts are available at https://doi.org/10.5281/zenodo.7881059.

**Table 2:**
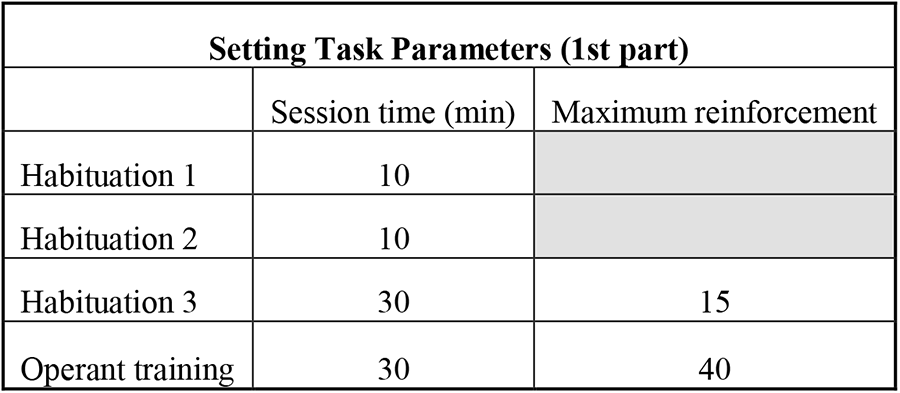
Task parameters used for the individual stages during the initial training. The depicted parameters follow the interface of the Med-PC V software (MedAssociates). They may slightly differ when adapted to a different type of software.

**Table 3:** Basic task parameters used for the individual stages during the advanced phase of the training. The depicted parameters follow the interface of the Med-PC V software (MedAssociates). They may slightly differ when adapted to a different type of software.

| Setting Task Parameters (2nd part) |  |  |  |  |  |  |  |  |  |  |
| --- | --- | --- | --- | --- | --- | --- | --- | --- | --- | --- |
|  | first correct lever<br>(1=left, 2=right) | first correct lever<br>(1=left, 2=right) | reinforcement device<br>(1=pellet, 2=dipper, 3=drug) | reinforcement time (sec) | time out following reinforcement (sec) | session time (min) | first fixed ratio value | second fixed ratio value | maximum reinforcement | softcr data array (1=yes, 0=no) |
| LR sequence | 1.000 | 0.000 | 1.000 | 0.050 | 0.000 | 60.00 | 2.0 | 1.0 | 40.000 | 1.000 |
| LLR sequence |  |  |  |  |  |  |  |  |  |  |
| LLRR sequence |  |  |  |  |  |  |  |  |  |  |
| LLRR with baseline time limits |  |  |  |  |  |  |  |  |  |  |
| LLRR with stricter time limits |  |  |  |  |  |  |  |  |  |  |
| LLRR alternating: 3xLLRR + 1x LLR |  |  |  |  |  |  |  |  |  |  |
| LLRR LEFT |  |  |  |  |  |  |  |  |  |  |
| LLRR Right |  |  |  |  |  |  |  |  |  |  |
| LLRR MIDDLE |  |  |  |  |  |  |  |  |  |  |
| Extinction |  |  |  |  |  |  |  |  |  |  |

**Table 4:**
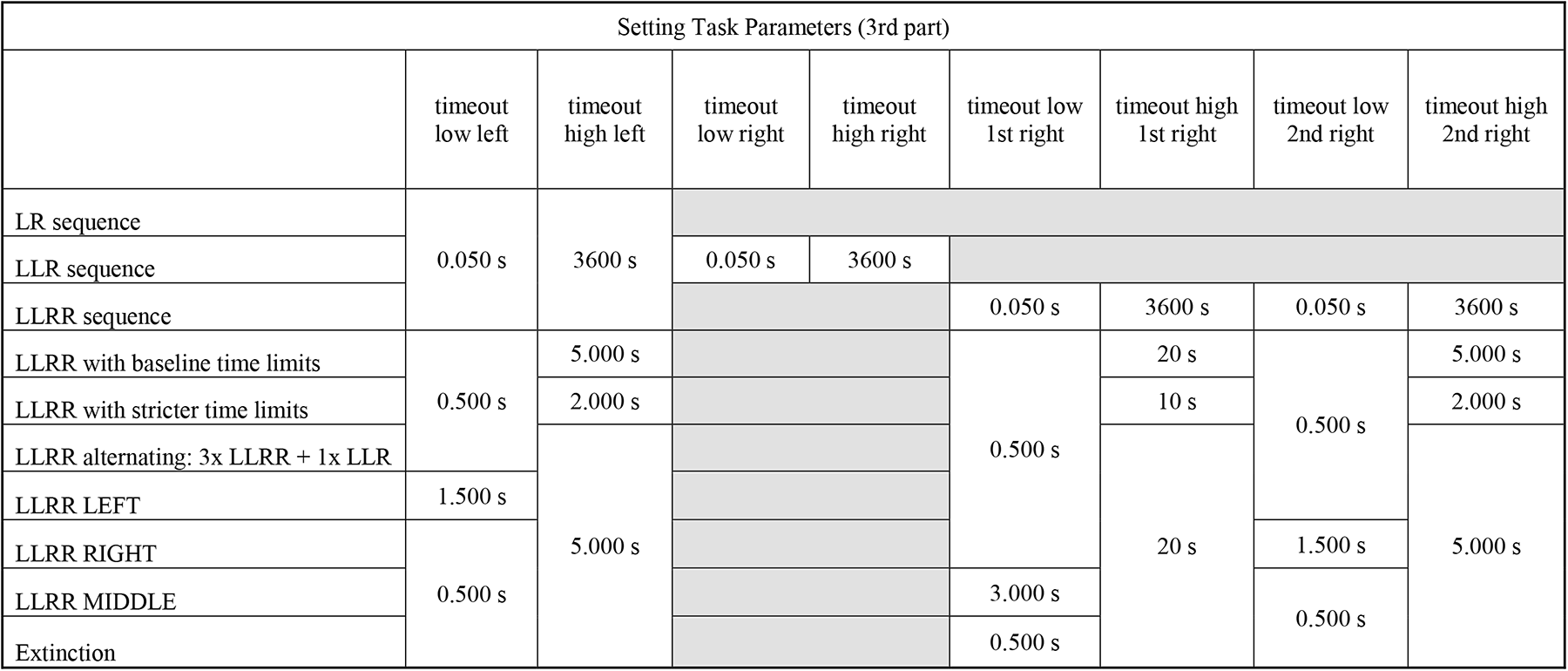
Timing task parameters used in advanced stages of the training. The depicted parameters follow the interface of the Med-PC V software (MedAssociates). They may slightly differ when adapted to a different type of software.

### Statistical analysis

We used 22 mice in total, 11 mice per group (control vs. VPA). Outliers were defined as values outside the range (average +/-2 x standard deviation) and they were excluded from the analysis. The identified outliers are indicated in figure legends together with respective sample sizes. Statistical analysis was performed with GraphPad Prism version 8 (GraphPad Software). We used two t-test and one-way ANOVA for evaluating the effect of one variable and two-way ANOVA or repeated measures (RM) two-way ANOVA for evaluating the effects of two variables. Sidak’s test was used for post-hoc comparison. For the analysis of the cumulative data we used a non-parametric Mann-Whitney test to compare individual data points between individual groups of animals. Significance was set at p < 0.05 and data in graphs are presented as average +/-SEM. Results of statistical analyses are reported as 95 % confidence intervals for the difference between means (95 % CI) and respective p values.

### Data and code accessibility

The code used to run the task is available at https://doi.org/10.5281/zenodo.7881104. The scripts for data analysis are available at https://doi.org/10.5281/zenodo.7881059. The raw data presented in the article are available at https://doi.org/10.5281/zenodo.7875357.

## Results

### Evaluation of the autism-like phenotype in the VPA mice

First, we used the social preference test, hole-board test and sticky tape test to confirm the mice prenatally exposed to VPA show the typical autism-like phenotype. Results indicating social and dexterity impairment are shown in Fig. 3.

**Figure 3:**
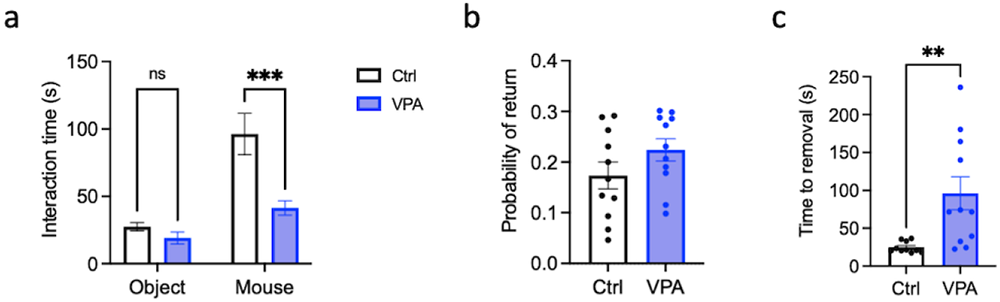
VPA mice show social impairment and reduced dexterity. (a) Interaction times during the social preference test in VPA and control mice. n(VPA, ctrl)=11. Effect of treatment, 95 % CI [14.4; 49.0], p=0.0007; effect of stimulus, 95 % CI [-62.9; -28.3], p<0.0001; two-way ANOVA. (b) Probability of returning to the same hole during the hole board test in VPA and control mice. n(ctrl, VPA)=11. 95 % CI [-0.02; 0.12], p=0.157; two-tailed t test. (c) Time to remove tape in VPA and control mice during the tape removal task. n(ctrl)=10 (1 outlier excluded), n(VPA)=11. 95 % CI [22.9; 1119.7], p=0.0061; two-tailed t test. All data are shown as mean ± SEM. **, p < 0.01; ***, p < 0.001.

### Both control and VPA mice can learn the timed sequence task and they improve during training

All mice met the habituation criteria and finished the subsequent training. The habituation was usually completed within one session, only in VPA group, one mouse 1x repeated Habituation 2 and 1x Habituation 3 session. There was no significant difference between the control and VPA mice in the number of sessions needed to reach the training criteria (Fig. 4a). Both groups also significantly improved their performance during training. We compared their performance in the 1st and the last session of the “LLRR baseline” timed paradigm, that is, in the 1st timed training session and in the last baseline session, just before the extinction. While mice were reaching the same number of rewards in both sessions (Fig. 4b), both control and VPA mice significantly decreased the number of incorrect presses and presses emitted during timeout (TO presses) (Fig. 4c, d). Initially, VPA mice performed less TO presses compared to controls but the difference was less prominent after training (Fig. 4e, f). To understand the mice activity during the session, we quantified individual types of pressing sequences, correct and incorrect, performed in the 1st and last LLRR baseline session. Unlike the 1st session, in the last session mice performed significantly more correct sequences than any other type of sequence (Fig. 4g, h). During sessions, both groups often alternated periods of activity (bouts) and inactivity (pauses). Most of the bouts and pauses were relatively short (less than 5 and 2 min, respectively) and their duration did not differ between control and VPA mice (Fig. 5a, b). We also analyzed intervals between the individual presses in the 1st and last session of the LLRR baseline paradigm, only in correctly executed sequences. Although the required intervals between the two left and the two right presses were identical, both control and VPA mice were significantly faster to emit the second right press, especially after training (Fig. 5c, d). Initially VPA mice showed longer intervals between presses compared to controls but during training, the average inter-press intervals in VPA and controls became indistinguishable (Fig. 5c, d). To assess the potential increase in stereotypic behaviour during training, we checked variation coefficients of individual inter-press intervals during the 1st and the last session of the LLRR baseline paradigm and we found decrease in the variability in the middle interval (L-R) in VPA animals only (Fig. 5e-j).

**Figure 4:**
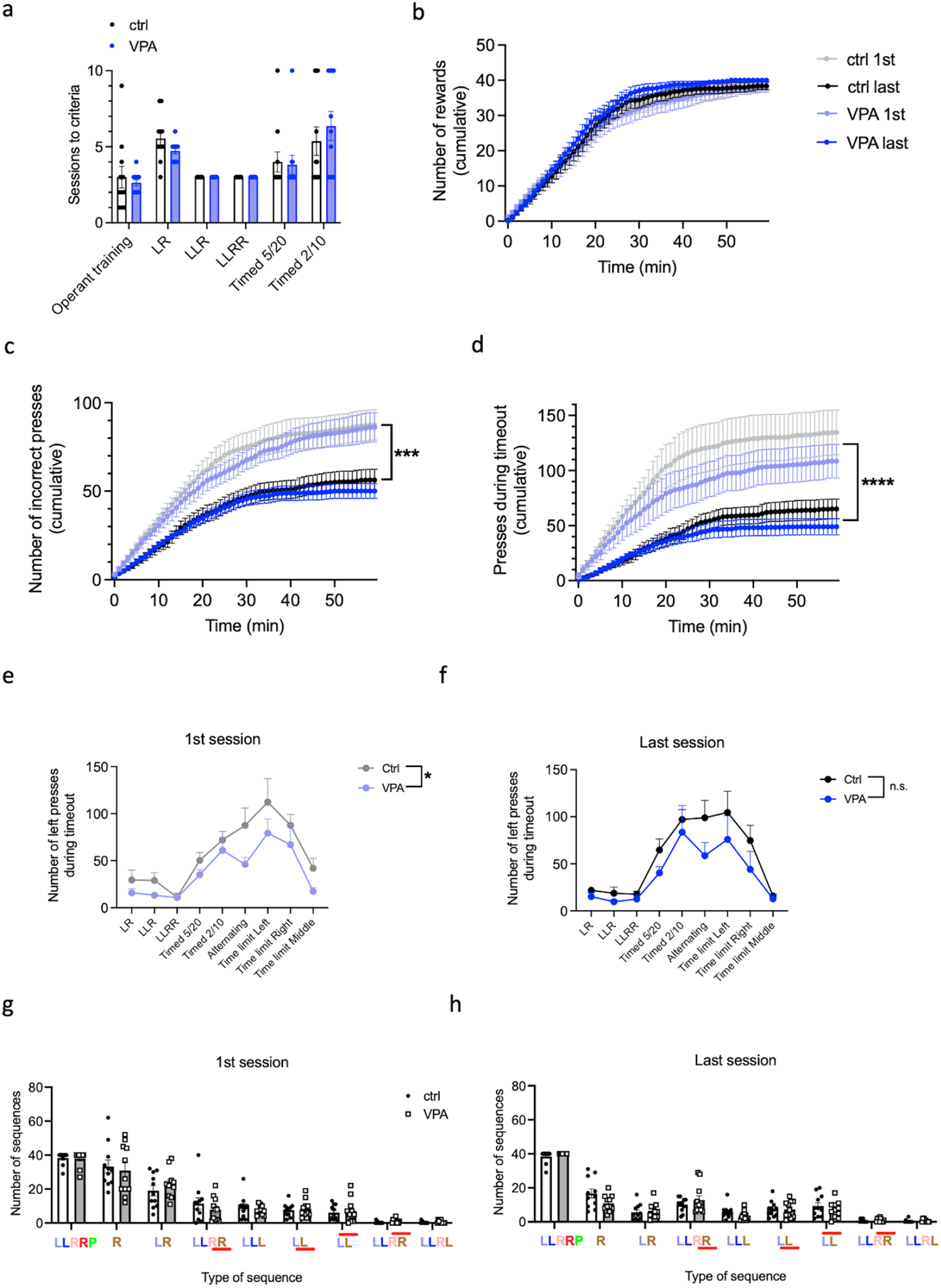
VPA and control mice learn successfully all stages of the timed sequence task. (a) Number of training sessions needed to reach performance criteria in individual training stages. (b-d) Cumulative graphs showing number of rewards earned (b), incorrect presses (c) and presses during timeout (d) emitted by VPA and control mice in the 1^st^ (light symbols) and last (dark symbols) session of the baseline timed LLRR sequence. (e, f) Number of left timeout presses emitted on the 1st (effect of treatment, 95 % CI [0.995; 38.0], p=0.0399) (e) and last (effect of treatment, 95 % CI [-6.41; 42.2], p=0.1403) (f) sessions at different stages of training. (g, h) Different types of pressing sequences performed on the 1st (g) and last (h) session of the baseline timed LLRR sequence. Sequence abbreviations: LLRRP – correct sequence followed by a reward (“Pellet”); R – sequence started incorrectly with Right press; LR – second press incorrect Right; LLR<u>R</u> - fourth press too fast (below threshold); LLL - third press incorrect Left; L<u>L</u> - second press too fast (below threshold); LL - second press too slow (above threshold); LLRR - fourth press too slow (above threshold); LLRL - fourth press incorrect Left. In all graphs, n(ctrl, VPA)=11. Data are shown as mean ± SEM and they were analyzed by two-way ANOVA (g, h), RM two-way ANOVA (a, e, f) or Mann-Whitney test (b-d). *, p<0.05; ***, p < 0.001; ****, p<0.0001.

**Figure 5:**
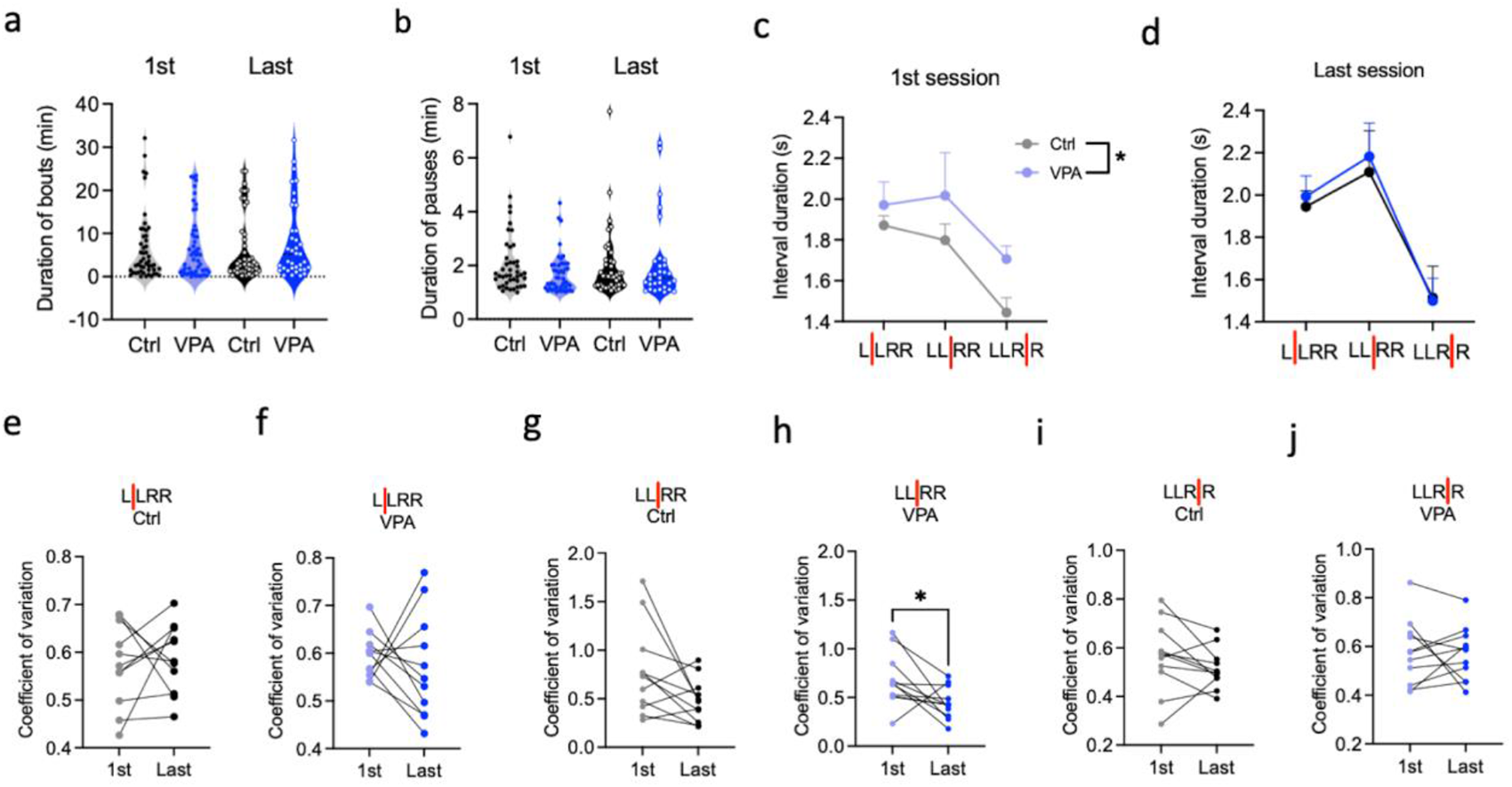
VPA mice show longer inter-press intervals and decreased variability during specific stages of training. (a-b) Duration of pressing bouts (a) and pauses (b) during the 1st and last session of the baseline timed LLRR sequence. (c-d) Duration of individual inter-press intervals during the 1st (effect of treatment, 95 % CI [-0.376; -0.011], p=0.0381; effect of interval, p=0.0037) (c) and last (effect of treatment, 95 % CI [-0.259; 0.187], p=0.750; effect of interval, p<0.0001) (d) session of the baseline timed LLRR sequence. (e-j) Comparison of the variability of inter-press intervals in VPA (f, h, j) (in h, 95 % CI [-0.452; -0.014], p=0.0391) and control (e, g, i) mice during the 1st and last session of the baseline timed LLRR sequence. In all graphs, n(ctrl, VPA)=11. Data are shown as mean ± SEM and they were analyzed by one- (a, b) or two-way (c, d) ANOVA or by paired t test (e-j). *, p < 0.05.

### VPA mice show better adaptation to changed pressing times but not changed pressing sequence

After the mice showed reliable performance in the LLRR baseline paradigm, we manipulated either the sequence structure or the required inter-press intervals. First, we implemented LLRR *alternating* paradigm where in every 4th trial the LLRR sequence was shortened to LLR (Fig. 6a-d). The occasional shortening of the pressing sequence affected control and VPA mice differently. During the 3 days of LLRR *alternating* paradigm, controls decreased the number of premature and increased the number of delayed second right presses while VPA mice did the opposite (Fig. 6c, d). After alternating the sequence, we changed the individual inter-press intervals, specifically by prolonging the lower time limit and forcing animals to wait longer before the next press. First, we increased the lower time limit between the two left presses to 1.5 s which had only modest effect on mice’s performance (data not shown). Next, we prolonged the lower time limit between the two right presses to 1.5 s. This affected mice’s performance (Fig. 6e-m) but unexpectedly, VPA mice adapted better to this change and during the three training days, they decreased the number of incorrect presses (Fig. 6g, i). In particular, VPA mice were less prone to make premature second right press (Fig. 6i, m). Finally, we changed the interval between the left and right middle presses by increasing the lower limit to 3 s which proved to be the most challenging modification. Although VPA mice again adapted easier to this change, the difference between groups was not so prominent (Fig. 7a-j). VPA mice did not make less premature first right presses compared to controls (Fig. 7i) but interestingly, they were less likely to change the sequence structure by adding 3rd left press (Fig. 7h). Overall, our data suggest that VPA mice can easier adapt to changes in the inter-press intervals but they are less likely to change the number and order of presses in the originally learned sequence.

**Figure 6:**
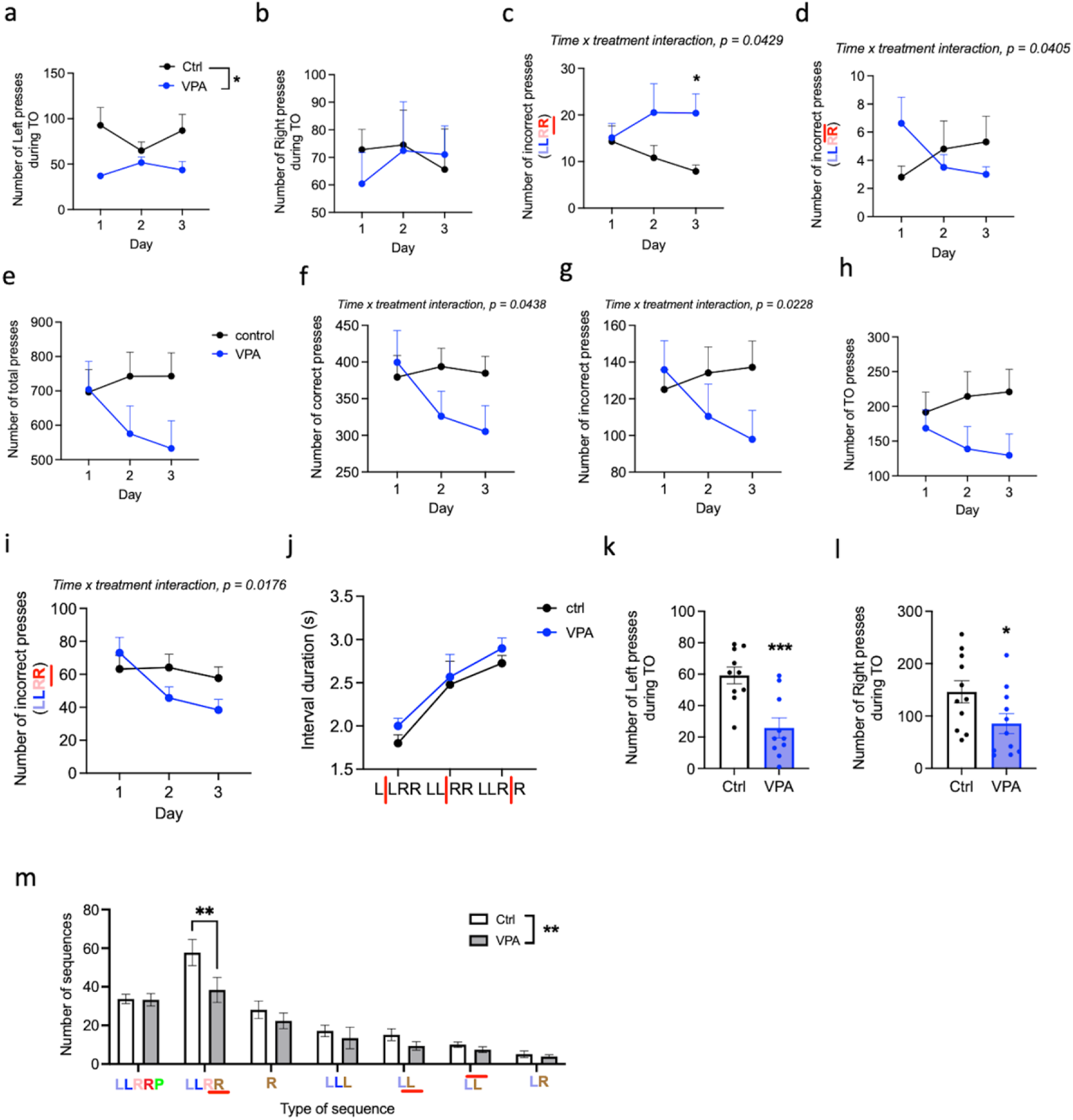
VPA mice show better adaptation to the change of timing but not when the pressing sequence itself is changed. (a-d) Performance criteria during the LLRR alternating sessions. Number of left (effect of treatment, 95 % CI [4.69; 70.10], p=0.027) (a) and right (b) presses during timeout and number of fourth presses emitted too fast (c) and too slow (d). n(ctrl)=10 (1 outlier excluded from a-d), n(VPA)=8-10 (2 outliers excluded from a-b, 1 excluded from c and 3 excluded from d). (e-i) Performance criteria during the three LLRR *RIGHT* sessions. Number of total (e), correct (f), incorrect (g) and timeout (h) presses in each session. (i) Number of the fourth presses emitted too fast in each session. n(ctrl, VPA)=11. (j-q) Performance criteria during the last LLRR *RIGHT* session only. (j) Duration of individual inter-press intervals. (k, l) Number of left (95 % CI [-50.9; -15.9], p=0.0008) (k) and right (95 % CI [-120.1; -1.163], p=0.046) (l) timeout presses. (m) Different types of pressing sequences emitted (effect of treatment, 95% CI [1.52; 9.57], p=0.007). Sequence abbreviations as in Fig. 4. n(ctrl)=10-11 (1 outlier excluded from k), n(VPA)=10-11 (1 outlier excluded from k). Data are shown as mean ± SEM and they were analyzed by RM two-way ANOVA (a-j), two-tailed t test (k, l) or two-way ANOVA (m). *, p < 0.05; **, p < 0.01; ***, p < 0.001.

**Figure 7:**
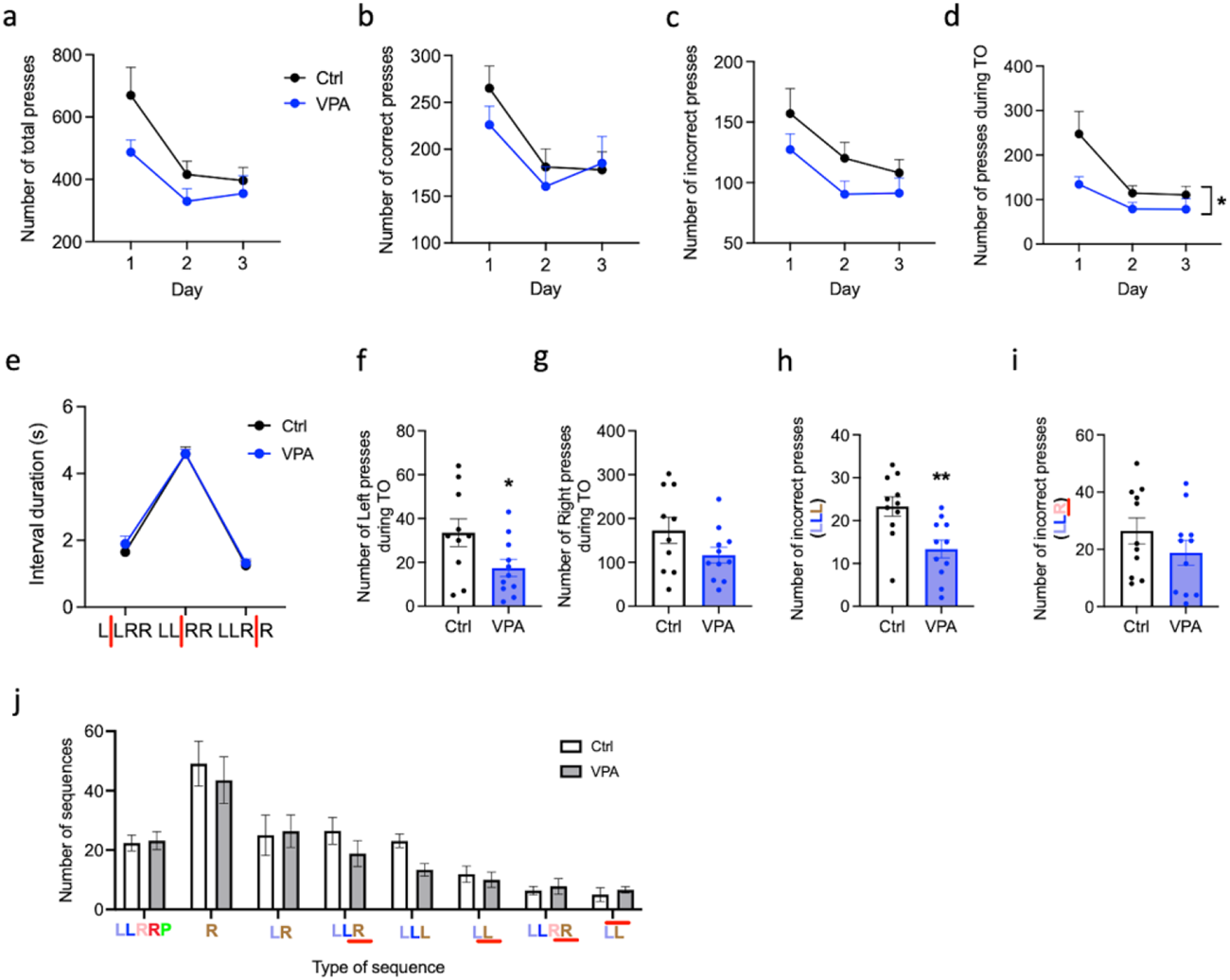
VPA mice maintain the original sequence when timing of the presses is changed. (a-d) Performance criteria during the three LLRR *MIDDLE* sessions. Number of total (a), correct (b), incorrect (c) and timeout (effect of treatment, 95 % CI [8.0; 112.9], p=0.026) (d) presses in each session. (e-n) Performance criteria during the 1st LLRR *MIDDLE* session only. (e) Duration of individual inter-press intervals. (f, g) Number of left (95 % CI [-31.30; -0.79], p=0.040) (f) and right (95 % CI [-127.7; 15.0], p=0.115) (g) timeout presses. (h) Number of third presses incorrect Left. 95 % CI [-16.36; -3.46], p=0.004). (i) Number of third presses emitted too fast. 95 % CI [-20.69; 5.42], p=0.237. (j) Different types of pressing sequences emitted during the 1st LLRR *MIDDLE* session. Sequence abbreviations as in Fig. 4. n(ctrl)=10-11 (1 outlier excluded from f, g), n(VPA)=11. Data are shown as mean ± SEM and they were analyzed by RM two-way ANOVA (a-d), two-way ANOVA (e, j) or two-tailed t test (f-i). *, p < 0.05; **, p < 0.01.

### VPA mice are less affected by the omission of reinforcement in the extinction session

Finally, we performed a single extinction session to test if VPA mice were more likely to maintain their performance in the absence of reward, which would be a sign of cognitive inflexibility (Mar et al., 2013). Control and VPA mice on average completed the same number of trials and a similar proportion of mice in both groups did not complete the maximum 40 trials (Fig. 8a, g). In line with that, the number of total and correct presses in both groups also did not differ (Fig. 8b, c, h). However, VPA mice emitted less incorrect and TO presses (Fig. 8d-f). They were less prone to start a new sequence with the right press, after the reward failed to be delivered in the end of the sequence (Fig. 8j). Instead, VPA mice responded by prolongation of the last inter-press interval (Fig. 8i), from approximately 1.5 s (Fig. 5d) during the baseline performance to almost 2 s (Fig. 8i). In summary, similar to the previous sessions, VPA mice tended to maintain the original order of presses in the sequence while having less difficulty to change the time intervals between them.

**Figure 8:**
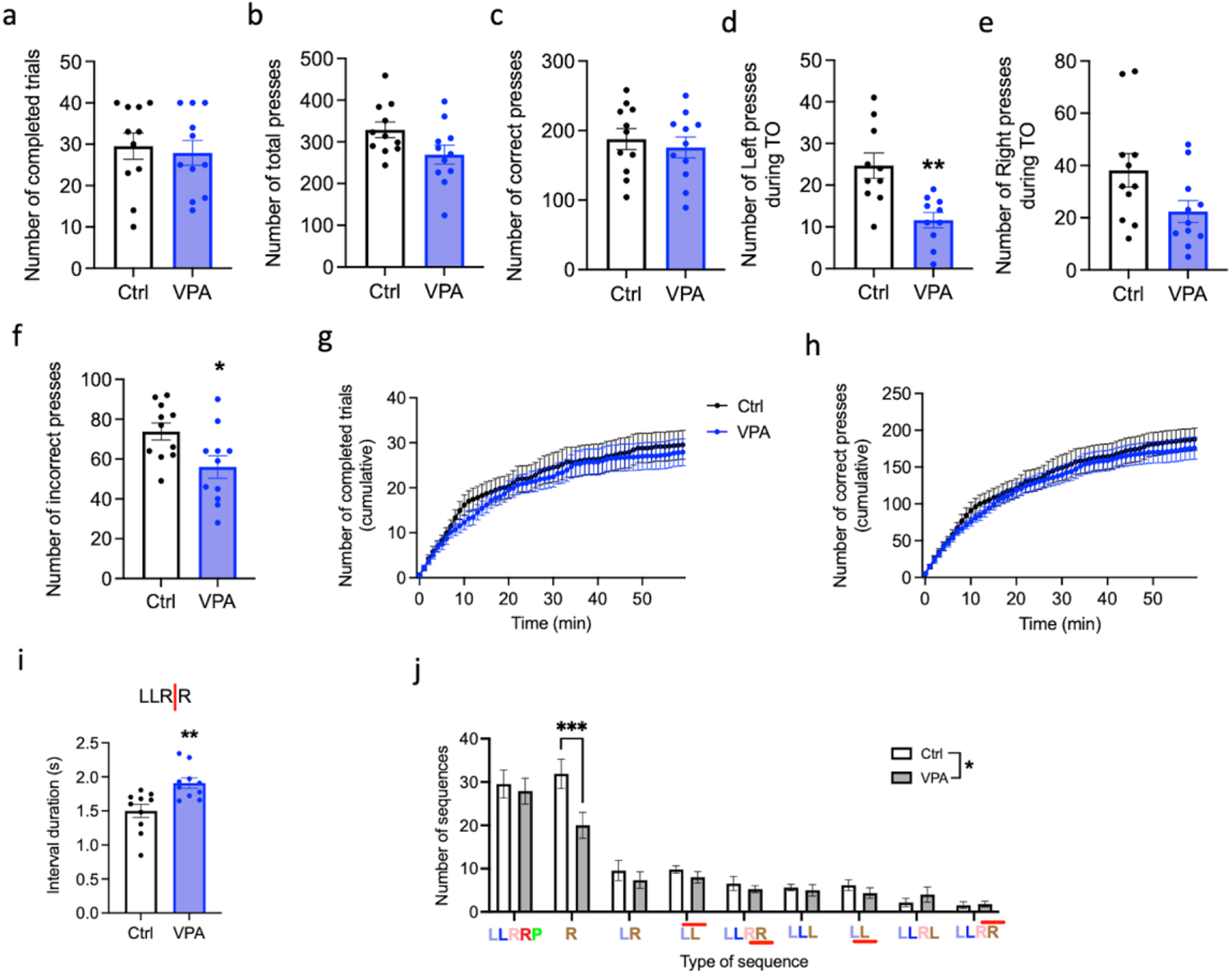
VPA mice maintain the original sequence when the reward is omitted during extinction session. (a-f) Performance parameters during extinction session. Number of completed trials (95 % CI [-10.75; 7.47], p=0.712) (a), total presses (95 % CI [-120.6; 2.10], p=0.058) (b), correct presses (95 % CI [-56.5; 32.3], p=0.576) (c), left (95 % CI [-20.57; -5.63], p=0.002) (d) and right (95 % CI [-31.83; 0.195], p=0.053) (e) presses during timeout and incorrect presses (95 % CI [-32.6; -3.1], p=0.021) (f). (g, h) Cumulative graphs showing number of completed trials (g) and correct presses (h). (i) Duration of the last inter-press interval. 95 % CI [0.151; 0.667], p=0.0038. (j) Different types of pressing sequences emitted during the Extinction session. Effect of treatment, 95 % CI [0.368; 3.895], p=0.018. Sequence abbreviations as in Fig. 4. n(ctrl)=10-11 (1 outlier excluded in d and i), n(VPA)=10-11 (1 outlier excluded in d and i). Data are shown as mean ± SEM and they were analyzed by two-tailed t test (a-f, i), Mann-Whitney test (g, h) or two-way ANOVA (j). *, p < 0.05; **, p < 0.01; ***, p < 0.001.

## Discussion

We present a new paradigm that allows evaluation of motor learning and flexibility. The design of the task was motivated by the need to test a complex behavior in an automated and reproducible manner, while using a commonly available equipment. In order to understand behavioral changes in various mouse models, it is necessary to evaluate multiple parameters. However, in the operant tasks, often only a few primary parameters such as accuracy are commonly analyzed which may lead to disregard of more subtle differences in behavioral strategies. For instance, we showed that in the VPA mice, the total number of earned rewards was indistinguishable from controls at all stages of the training. Upon closer look however, the performance of the two groups was quite different, suggesting slower motor performance and decreased ability to adapt the originally learned sequence. Another significant difference between the VPA and control mice was the increased number of TO presses present at multiple stages of the training (Fig. 4e).

In our task, we take advantage of gradually increasing difficulty at different stages to reveal potentially gross or more subtle behavioral impairment and we analyze multiple parameters to better understand its character. During the training, mice have to employ basic operant learning, motor learning, time perception, habit formation, behavioral flexibility and motivation. Importantly, even with only two fixed levers, the task consists of reward-distal and reward-proximal elements and allows to analyze them separately (Keeler et al., 2014). Although the two left and the two right presses were seemingly symmetrical and the required pressing intervals between them were identical at baseline, mice were clearly able to distinguish their distinct reward proximity as the average inter-press interval between the two right presses was significantly shorter (Fig. 5d). In order to use the timed sequence task for the evaluation of flexibility, we need to make sure the mice performance is becoming automated and habitual during the training. To test this, we compared the variability of individual pressing intervals in the 1st and the last “LLRR baseline” session (Fig. 5e-j). Surprisingly, we did not find any decrease of variability in control animals, possibly because the pressing intervals were largely constricted by the pre-defined criteria and there was little space for further improvement. In addition to the low variability of individual motor actions, the automated character of behavior is indicated by an increase in speed and accuracy (Turner et al., 2022). In our experiment, we observed a significant increase in accuracy (Fig. 4c) and efficiency, as mice performed markedly less TO presses in later stages of the training (Fig. 4d).

There are currently several alternatives that allow a detailed testing of motor or operant learning and flexibility. Among those, the touchscreen-based systems offer a great number of pre-designed paradigms and allows creating additional, custom made tasks (Horner et al., 2013; Mar et al., 2013; Heath et al., 2016; Janickova et al., 2021). However, a significant cost of such systems may be prohibitive for some users. Standard operant boxes, on the other hand, are easily accessible to most laboratories. We show here that even the most basic operant box equipped with two fixed levers and one feeder can be used to perform a relatively complex task and collect detailed behavioral data. We provide a computer code for the task itself and also a set of scripts for data analysis, allowing easy and fast processing of large data sets. In mice we tested, completing of the whole paradigm took approximately 45 training days. This is more than in standard operant learning tasks but comparable to other more sophisticated paradigms offering detailed insight into mice behavior. One of the advantages of our paradigm is that it is testing the animals’ performance over the course of several weeks at different levels of difficulty and it records and analyzes multiple behavioral parameters. It makes it perfectly suited to detect more subtle changes in behavior, compared to one-session tasks such as hole board test. Due to its long-term character, it can also determine if specific behavioral changes can manifest temporarily at the beginning of training or if they are more consistent, suggesting a permanent cognitive trait.

A potential limitation of the study is our focus on examining only males. We decided to validate our paradigm in males due to the higher prevalence of ASD in men (Werling and Geschwind, 2013). In addition, most studies did not find a difference in the behavioral phenotype between sexes after prenatal VPA exposure (Ergaz et al., 2016). However, conducting the paradigm on females would be intriguing since they can exhibit lower cognitive flexibility than males (Korol et al., 2004; LaClair et al., 2019). Besides sex, another variable to consider is the influence of circadian rhythm. Mice typically exhibit higher activity during nocturnal periods. Additionally, VPA treated C57BL/6 mice displayed decreased place re-learning and an increased tendency to perseveration, particularly during light periods (Puścian et al., 2014). These findings align with observations that people with ASD often display nocturnal activity and during the day, they have increased melatonin levels, which contribute to emotional or cognitive problems (Carmassi et al., 2019).

In summary, we provide a new easily applicable paradigm for detailed testing of motor learning and flexibility. We provide tools for data analysis, arguing that multiple parameters should be used to evaluate the animals’ performance in a complex task. Finally, we provide an example showing that our paradigm can reveal subtle differences in mice performance and help to identify general behavioral mechanisms underlying those differences.

